# Transcriptional heterogeneity of stemness phenotypes in the ovarian epithelium

**DOI:** 10.1101/2020.06.10.145045

**Authors:** LE. Carter, DP. Cook, CW. McCloskey, T. Dang, O. Collins, LF. Gamwell, HA. Dempster, BC. Vanderhyden

## Abstract

The ovarian surface epithelium (OSE) is a monolayer of epithelial cells covering the surface of the ovary. During ovulation, the OSE is ruptured to allow release of the oocyte. This wound is quickly repaired, but mechanisms of this repair are poorly understood. The contribution of tissue-resident stem cells in the homeostasis of several epithelial tissues is widely accepted, such as the intestinal epithelium, airway epithelium, and skin, but their involvement in OSE maintenance is unclear. While putative stem cell populations in the OSE have been described, how they are regulated is poorly defined. We show that traits associated with stem cells (stemness) can be increased in OSE following exposure to the cytokine TGFB1, overexpression of the transcription factor *Snai1*, or deletion of *Brca1*. By assessing the gene expression profiles of these populations, we show that stemness is often linked to mesenchymal-associated gene expression and higher activation of ERK signalling, but it is not consistently dependent on their activation. Expression profiles of these populations are extremely context specific, suggesting that stemness may not correspond to a single, distinct population, but rather is a heterogenous state that can possibly emerge from diverse environmental cues. Together, these findings support that the OSE may not require distinct stem cell populations for long-term maintenance, and may achieve this through transient dedifferentiation into a stem-like state.

## Introduction

It is thought that stem cell populations are responsible for long-term maintenance of many adult tissues. The characterization of stem cells associated with epithelial tissues has been an active field of research for the past few decades. While several distinct stem cell populations have been described, such as an LGR5+ population at the base of intestinal crypts^1^, it is unclear if all epithelial tissues are maintained by such a defined population. For example, it has been shown that following stem cell depletion, differentiated airway epithelial cells can dedifferentiate and become functional multipotent stem cells^2^. It is also unclear if stem cells are necessarily required to maintain epithelial tissues comprising a single cell type, as some baseline capacity for proliferation could maintain the entire tissue. In the mesothelium, for example, there have been reports of putative stem/progenitor cells for over two decades, but a well-defined stem cell population has yet to be identified^3^.

The ovarian surface epithelium (OSE) is a promising tissue for studying stemness dynamics in tissue maintenance. It is a monolayer of cells surrounding the ovary and, during each ovulation, this tissue is ruptured to facilitate release of an oocyte. Afterwards, the OSE layer is rapidly repaired^4–7^. Post-ovulatory wound repair is a poorly understood process despite ovulation being a major non-hereditary risk factor for ovarian cancer.

Several putative OSE stem cell populations have been described, each defined by different cell surface markers, including ALDH1A1, LGR5, LY6A (Sca-1), and more^8–11^. The relationships between populations described in these studies are still unclear. Further, it is unclear if these populations are static or can emerge from differentiated OSE in response to ovarian dynamics. We have previously shown that, like the mammary epithelium^12^, induction of an epithelial-to-mesenchymal transition (EMT) can transiently promote features of stem cells (stemness) in differentiated OSE^10^. This is particularly relevant in the context of ovulation, as the EMT is thought to be an important component of wound repair and the EMT-promoting cytokine TGFB1 is present in follicular fluid, bathing adjacent OSE at ovulation^10,13^. It is also secreted by macrophages at the ovulatory wound and granulosa cells during follicular development^14,15^.

Here, we further demonstrate that features of stemness can be enhanced in OSE cells. Profiling gene expression of different populations with enhanced stemness, we demonstrate that some features are relatively common, including EMT-associated expression patterns and higher activity of ERK and NFkB signalling, but global expression profiles are widely variable and stemness isn’t exclusively dependent on these common features. Together, this work supports that OSE tissue maintenance may not require a distinct stem cell population, but can emerge in response to their environment. Further, different environmental conditions can give rise to stem-like populations with different characteristics.

## Results

### CD44 is a marker of EMT-associated stemness in OSE cells

We have previously demonstrated that mouse OSE (mOSE) cells undergo an EMT and acquire stem cell characteristics when exposed to TGFB1^10,16^. To confirm these findings, we first assessed the ability of TGFB1-treated mOSE cells to form self-renewing spheroids in suspension culture. Treated cells formed over twice as many primary spheroids and, when dissociated and cultured, were more efficient at successfully generating secondary spheroids, confirming their ability to self-renew (**Fig. 1a**). Morphologically, mOSE spheres were large and compact, regardless of whether they had been treated (**Fig. 1b**). To validate this enhanced stemness in human cells, we performed these experiments on primary cultures of human OSE (hOSE) cells. These cultures have a low proliferation rate *in vitro*, so to minimize the impact of aggregation in our quantifications, we used methylcellulose-based suspension culture to immobilize the cells. While these conditions, along with the slower proliferation, resulted in smaller spheroids, TGFB1-treated hOSE cells formed 3 times as many spheroids as untreated cells (**Fig 1c**).

**Figure 1.**
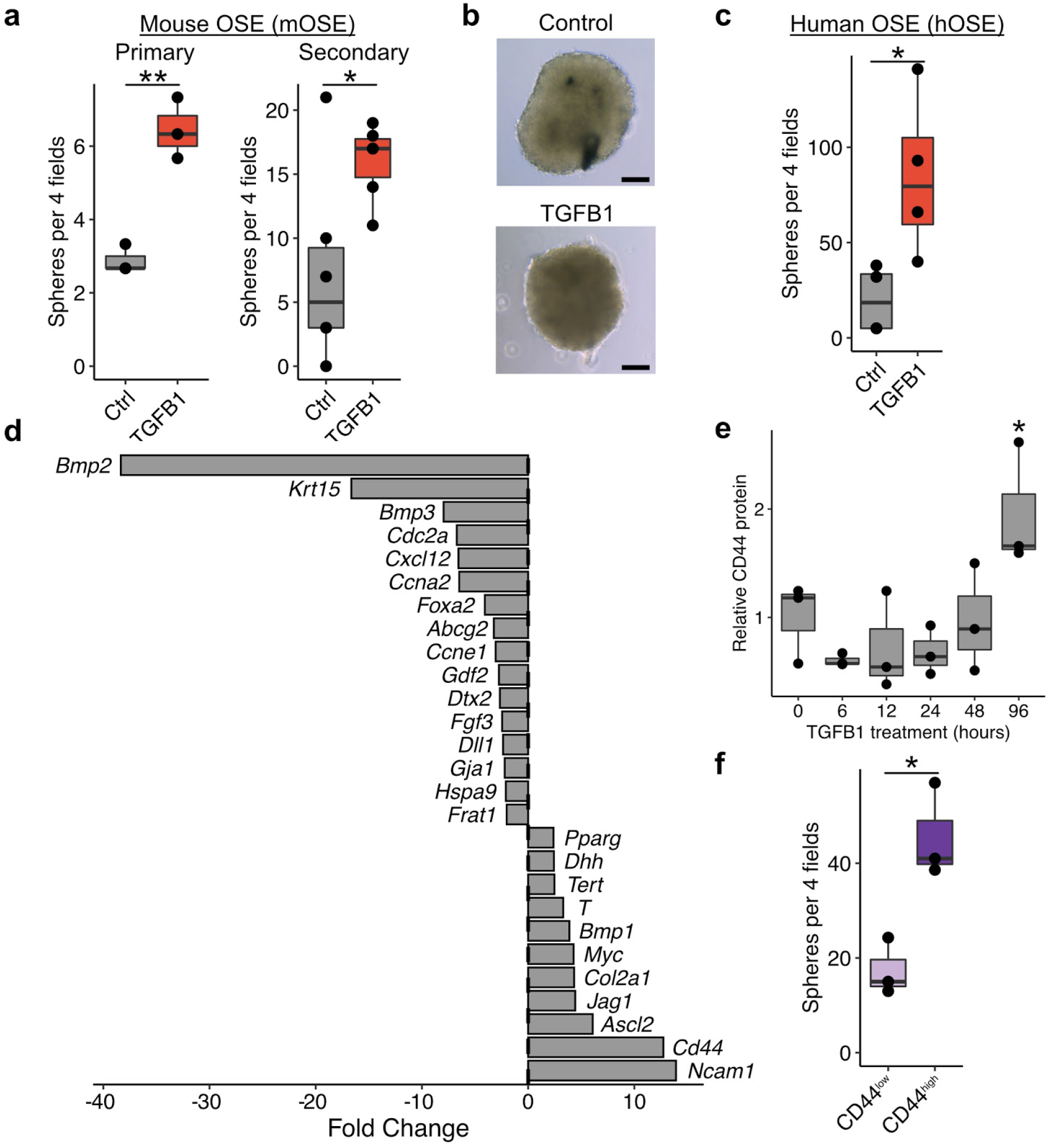
TGFB1 promotes stemness in the OSE. **a.** Primary (left) and secondary (right) sphere-forming capacity of mOSE cells treated with TGFB1 (10ng/mL). Data points represent the average number of spheres per 4 fields of view for each replicate. **b.** Phase contrast images of control and TGFB1-treated spheroids. Scale bar=100μm. **c.** Primary sphere-forming capacity of human OSE treated with TGFB1. **d.** Fold change values for a panel of putative stem cell markers in mOSE treated with TGFB1 for 4 days. **e.** Relative protein quantifications of CD44 throughout a time course of TGFB1 treatment in mOSE. Quantifications represent western blot pixel densitometry, normalized to B-actin and scaled to the mean intensity in untreated samples. A representative blot is included in Supplemental Figure 2b. **f.** Primary sphere-forming capacity of CD44- and CD44+ mOSE cells. All boxplots show median value (horizontal black line), estimated 25th and 75th percentiles, and whiskers represent 1.5 times the interquartile range. Linear regression models were used for all statistical tests. * *p*<0.05, ** *p*<0.01.

To determine if this enhanced stemness is associated with the expression of previously reported markers of OSE stem cells, we measured their expression throughout TGFB1 treatment in mOSE cells. *Aldh1a1, Lgr5*, and *Nanog* did not increase throughout a week of treatment, and in some cases even decreased over time (**Supplementary Fig. 1**). While this does not preclude the possibility of these genes being valid markers of stem cell populations *in vivo*, these results suggest that their regulation is independent from TGFB1-associated stemness.

To identify putative markers that are associated with this stemness, we assessed the expression of a larger panel of markers from a commercial “Stem Cell Marker” qPCR array (**Fig. 1d**). This identified several highly upregulated markers following 7 days of TGFB1 treatment, including *Ncam1* (14-fold), *Cd44* (13-fold), and *Ascl2* (6-fold) (**Fig. 1d**). CD44 has long been associated with stemness in mammary epithelial cells^17^ and more recently in the oviductal epithelium^18^. To determine if CD44 can be used as a selective marker to enrich for stem cell characteristics, we first validated that TGFB1 increases CD44 protein levels in mOSE cells (**Fig. 1e; Supplementary Fig. 2**) and then sorted CD44^high^ cells from TGFB1-treated mOSE by fluorescence-activated cell sorting (FACS). When placed in suspension culture, CD44^high^ cells formed approximately 2.5 times as many spheroids (**Fig. 1f**).

### Transcriptional profiling of mOSE stemness

We next sought to define a global profile of stemness, beyond a small number of markers. Spheroids themselves have been demonstrated to be enriched with stem/progenitor populations^19–21^. Since untreated mOSE cultures are capable of sphere formation, albeit at a lower frequency than TGFB1-treated mOSE cells, we reasoned that the transcriptional profile of these spheroids may represent an intrinsic stemness program, independent from exogenous factors. To compare this with TGFB1-induced stemness, we performed RNA-seq on mOSE cells cultured as a monolayer or as spheroids, each with and without TGFB1 treatment.

Untreated mOSE cells cultured as spheroids exhibited striking differences from those cultured in a monolayer, with 4950 differentially expressed genes between the conditions (*p* < 0.05, absolute log fold change > 0.5) (**Fig. 2a; Supplemental Data 1**). Using an aggregate reference of GO terms, KEGG pathways, Reactome pathways, and MSigDB Hallmark gene sets from the Molecular Signatures Database (MSigDB)^22,23^, we used gene set enrichment analysis (GSEA) to identify biological features associated with these changes (**Fig. 2c; Supplemental Data 2**). Spheroids were associated with decreased cell cycle, epithelial cell adhesion and, interestingly, DNA repair. Along with these changes, spheroid culture activated expression of chemokine signalling and wound repair programs. We also note that CD44, which we had used as a selection marker for stemness in TGFB1-treated mOSE, was also expressed over 4-fold higher in spheroids, whereas *Aldh1a1, Lgr5*, and *Ly6a* (*Sca-1*) were unchanged (**Fig. 2b**). We next used the PROGENy algorithm to infer changes in signalling pathway activity across these samples that may be contributing to these differences. Spheroids were associated with increased activity of many signalling pathways, with the largest increases in Hypoxia, NFkB, and MAPK signalling (**Fig 2d**). While GSEA results suggest several EMT-related changes, TGFB1 and WNT signalling are interestingly reduced, suggesting that this EMT program may be activated through NFkB or ERK (**Fig 2d**).

**Figure 2.**
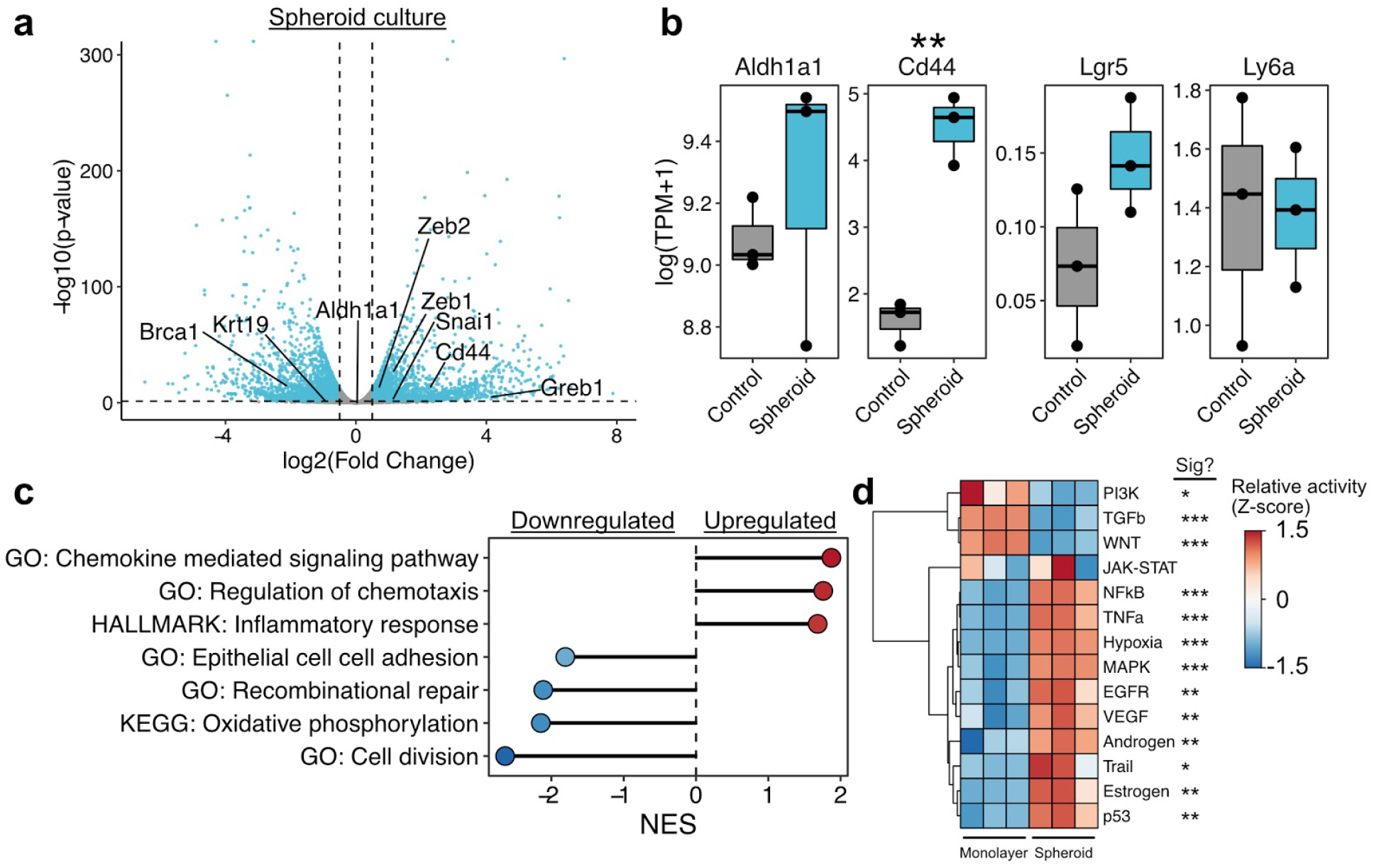
Transcriptional profile of intrinsic mOSE stemness. **a.** Plot showing the distribution of differentially expressed genes in mOSE cells cultured as spheroids relative to a monolayer. Each point corresponds to a single gene. Selected genes related to stemness and/or the EMT are highlighted on the plot. Dashed lines correspond to significance criteria (absolute log fold change > 0.5, *p* <0.05). **b.** Boxplots showing the expression values of putative mOSE stemness markers in mOSE cells cultured in a monolayer (Control) or as spheroids. Boxplots show median value (horizontal black line), estimated 25th and 75th percentiles, and whiskers represent 1.5 times the interquartile range **c.** GSEA results for selected gene sets enriched in differentially expressed genes in mOSE spheroids. All gene sets are significantly enriched (*p*<0.05) and normalized enrichment scores (NES) are shown. **d.** Inferred pathway activity in monolayer- and spheroid-cultured mOSE cells. Linear models were used for statistical testing for (b) and (d). * *p*<0.05, ** *p*<0.01, *** *p*<0.001.

While TGFB1 signalling was decreased in untreated spheroids, exogenous TGFB1 treatment enhanced stemness, increasing the proportion of cells capable of forming self-renewing spheroids. While these results are seemingly contradictory, week-long exposure to exogenous TGFB1 may activate similar expression programs through secondary effects or signalling crosstalk^24^. We next assessed expression changes associated with TGFB1 exposure. Week-long treatment of monolayer cultures resulted in 1508 differentially expressed genes (*p* < 0.05, absolute log fold change > 0.5) (**Fig. 3a; Supplemental Data 3**). This involved the activation of EMT-associated gene sets as expected, as well as a reduction in oxidative phosphorylation (**Fig. 3b; Supplemental Data 4**). While TGFB1 signalling was the only pathway inferred to have significantly altered activity, the estimated activity of EGFR and MAPK was higher in TGFB1-treated cells (*p* = 0.1 and 0.06, respectively) (**Fig. 3c**). Consistent with this, the GO term “ERK1 and ERK2 signalling cascade” was significantly enriched in upregulated genes following TGFB1 treatment (**Fig. 3b**).

**Figure 3.**
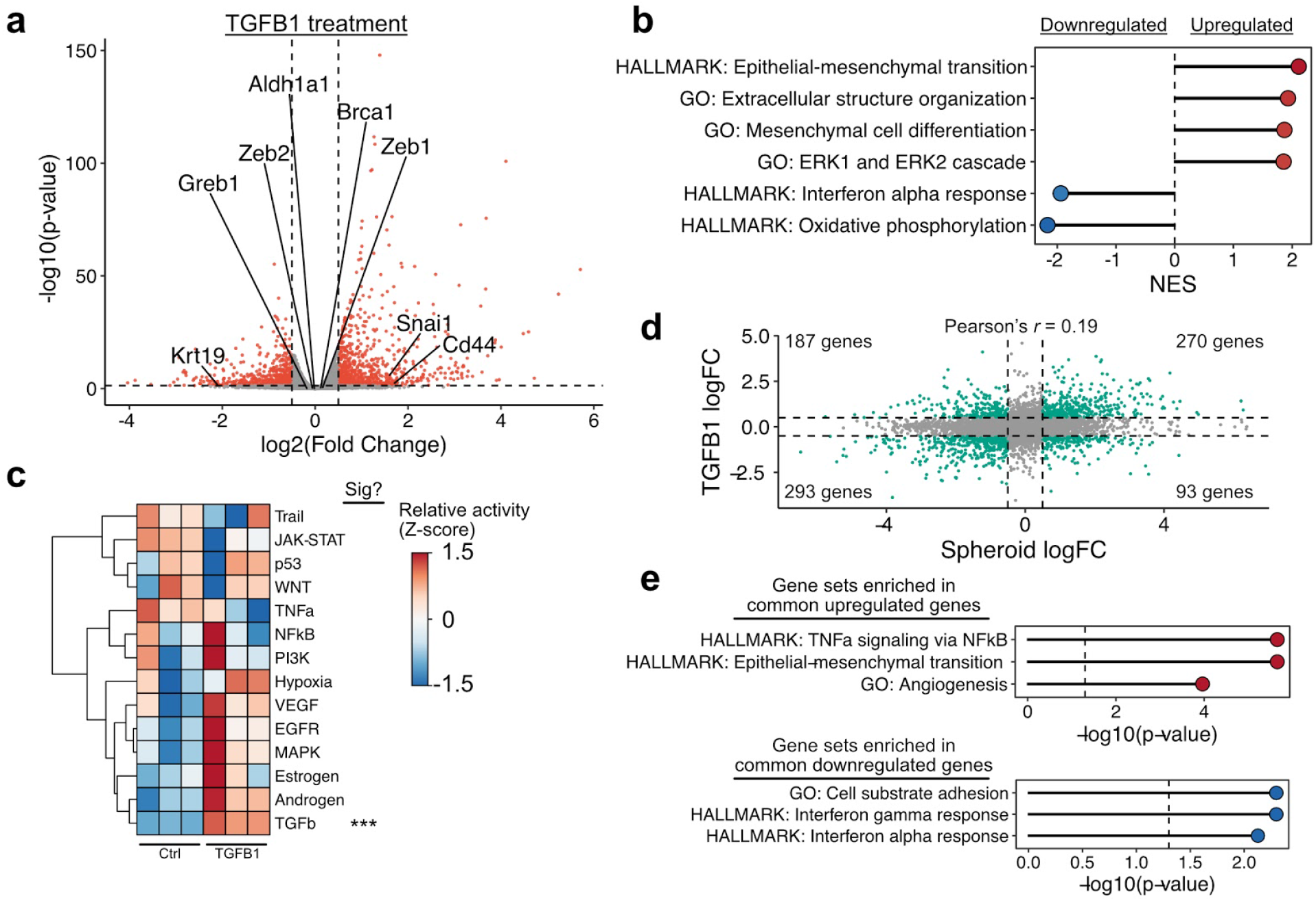
TGFB1 promotes a distinct stemness phenotype. **a.** Plot showing the distribution of differentially expressed genes in monolayers of mOSE cells treated with TGFB1 compared to untreated samples. Each point corresponds to a single gene. Selected genes related to stemness and/or the EMT are highlighted on the plot. Dashed lines correspond to significance criteria (absolute log fold change > 0.5, *p* <0.05). **b.** GSEA results for selected gene sets enriched in differentially expressed genes following TGFB1 treatment. All gene sets are significantly enriched (*p* <0.05) and NES values are shown. **c.** Inferred pathway activity in untreated and TGFB1-treated mOSE cells. *P*-values were computed from the *t* statistic of a linear regression model and were adjusted using the Benjamini-Hochberg false discovery rate (FDR) method. **d.** Plot comparing log fold-change values for spheroid-cultured and TGFB1-treated mOSE. Dashed lines correspond to fold change cutoffs used to assess significance. **e.** Plots showing gene sets enriched in commonly up- or downregulated genes following both TGFB1 treatment and spheroid culture. *P*-values were calculated using a Fisher exact test and were adjusted using the Bejamini-Hochberg FDR method.

This suggests that TGFB1 treatment initiates sequential or parallel signals similar to those present in spheroids. Consistent with this, untreated spheroids and TGFB1-treated mOSE monolayers have a significant overlap in expression changes relative to untreated mOSE cells cultured as a monolayer, sharing 270 upregulated genes and 293 downregulated genes (Fisher exact *p* = 3.0e-66 and 2.8e-117, respectively) (**Fig. 3d**). Conserved upregulated genes were strongly enriched for EMT-associated genes, NFkB signalling, and angiogenesis (**Fig. 3e**). Interestingly, very few gene sets were enriched in the conserved downregulated genes, with only interferon response and substrate adhesion genes being enriched (**Fig. 3e**). Together, this suggests that OSE stemness may involve higher activities of ERK and NFkB signalling and an associated reduction in typical epithelial traits.

### Snail activation promotes a unique stemness program in mOSE cells

Signalling pathways are highly pleiotropic and it is unclear if TGFB1-enhanced stemness is activated from core EMT regulatory networks or alternative components regulated by TGFB1. The EMT transcription factor *Snai1* (Snail) was upregulated in both TGFB1-treated mOSE cells and spheroids, so to test if EMT activation without exogenous cytokines could promote stemness, we derived mOSE cell lines with doxycycline-inducible Snail expression. Following Snail induction, cells had a higher sphere forming capacity than cells without doxycycline exposure, generating up to twice as many primary spheres and 3 times as many secondary spheres when passaged (**Fig. 4a**).

**Figure 4.**
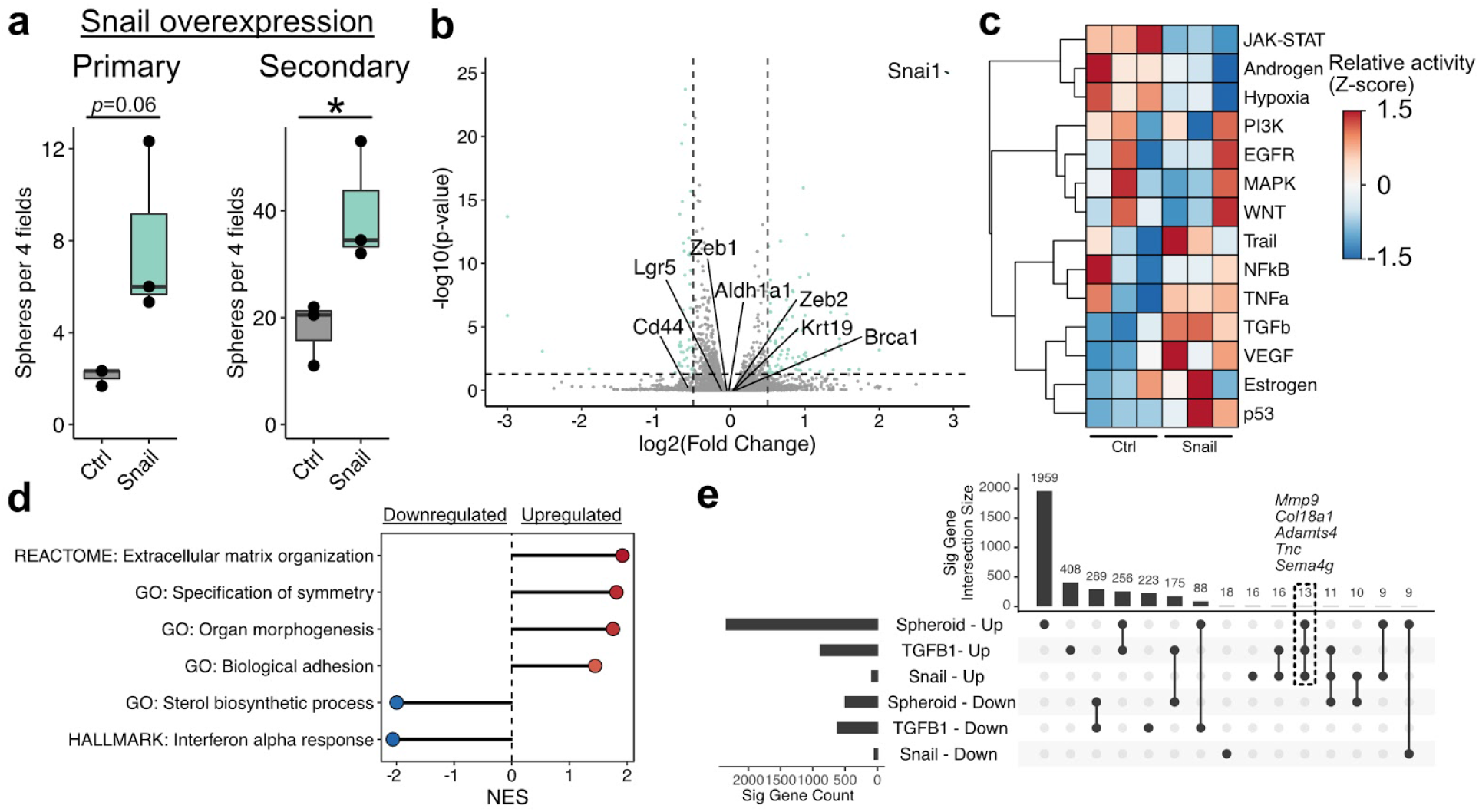
Snail overexpression promotes stemness with minimal gene expression changes. **a.** Primary (left) and secondary (right) sphere forming capacity of mOSE cells overexpressing Snail. Data points represent the average number of spheres per 4 fields of view for each replicate. Boxplots show median value (horizontal black line), estimated 25th and 75th percentiles, and whiskers represent 1.5 times the interquartile range. **b.** Plot showing the distribution of differentially expressed genes following Snail overexpression. Each point corresponds to a single gene. Selected genes related to stemness and/or the EMT are highlighted on the plot. Dashed lines correspond to significance criteria (absolute log fold change > 0.5, *p*<0.05). **c.** Inferred pathway activity in control and Snail-overexpressing mOSE cells. *P*-values were computed from the *t* statistic of a linear regression model and were adjusted using the Benjamini-Hochberg FDR method. No pathway is significantly different between conditions. **d.** GSEA results for selected gene sets enriched in differentially expressed genes following Snail overexpression. All gene sets are significantly enriched (*p*<0.05) and NES values are shown. **e.** UpSet plot showing overlaps in differentially expressed genes between all conditions assessed ranked by the condition/overlap with the largest number of genes. The top chart shows the intersection size for the conditions highlighted in the middle grid. A single, unconnected point corresponds to genes unique to only that condition. The total number of differentially expressed genes in each condition is shown in the left chart.

We next assessed the expression of the putative stem cell markers *Cd44* and *Sca-1*, which are both increased with TGFB1 treatment, and found that Snail induction had no effect on their expression, suggesting that their validity as markers of stemness may be context specific. To determine if Snail induction activates similar expression patterns to TGFB1-treated mOSE and spheroids, including higher ERK and NFkB activity, we performed RNA-seq on these cells with and without doxycycline. Snail-induced changes were more modest than with TGFB1 treatment or in spheroid culture, with only 85 upregulated and 44 downregulated genes (*p* < 0.05, absolute log fold change > 0.5; **Fig. 4b; Supplemental Data 5**). Interestingly, inferred pathway activity scores associated with EGFR, NFkB, MAPK, and TGFB1 were all unchanged following Snail induction (**Fig. 4c**). Further, no relevant gene sets associated with these pathways were enriched in the differentially expressed genes (**Fig. 4d**). The only gene sets associated with upregulated genes were largely related to cell morphology and extracellular matrix (ECM) remodelling (**Fig. 4d; Supplemental Data 6**). Consistent with TGFB1-treated mOSE and spheroids, the MSigDB Hallmark “Interferon Alpha Response” was the only gene set enriched in the downregulated genes following Snail induction (**Fig. 4d**). We note that of the 85 upregulated genes following Snail induction, 13 are shared with those commonly regulated in TGFB1 treatment and spheroids (**Fig. 4e**). These genes largely represent components of the ECM, including *Col18a1* and the metalloproteinases *Mmp9* and *Adamts4*. As the conditions share no consistently activated downstream signal that could be induced by ECM changes, these findings suggest that expression programs associated with stemness phenotypes are heterogeneous. Given frequent enrichment of gene sets associated with a mesenchymal phenotype, stemness may consistently involve higher levels of these traits, which can emerge from variable expression patterns^25^.

### BRCA1 loss promotes EMT-independent stemness in mOSE

Spheroids were associated with higher expression of many genes that were not affected by TGFB1. We noted that among these were several changes typically associated with ovarian cancer, including activation of the transcription factor *Pax8*, which is present in approximately 80% of ovarian tumors but not typically expressed in the OSE^26^; activation of *Greb1*, which promotes ovarian cancer growth^27^; and loss of *Brca1*, which, along with *Brca2*, is mutated in approximately 22% of high-grade serous ovarian tumours. Interestingly, loss of BRCA1 has been associated with promoting dedifferentiation and activation of EMT expression patterns in mammary epithelial cells^28^.

As the association between BRCA1 loss and stemness in the OSE had not been assessed, we next derived a primary mOSE line from *Brca1^tm1Brn^* mice harboring floxed *Brca1* alleles. To determine if BRCA1 loss enhanced stemness in these cells, we infected the cells with adenovirus containing either Cre recombinase (Ad-Cre) or GFP (Ad-GFP) as a control. Cre delivery, while not perfectly efficient, resulted in an approximately 60% reduction in BRCA1 levels across the population (**Supplemental Fig. 3**). When placed in suspension culture, cells with reduced BRCA1 formed over 5 times as many primary spheres and 3 times as many secondary spheres than control mOSE cells, suggesting that BRCA1 loss also enhances stemness in mOSE cells (**Fig. 5a**). These findings also support our previous finding that the expression of putative stemness markers may be context specific.

**Figure 5.**
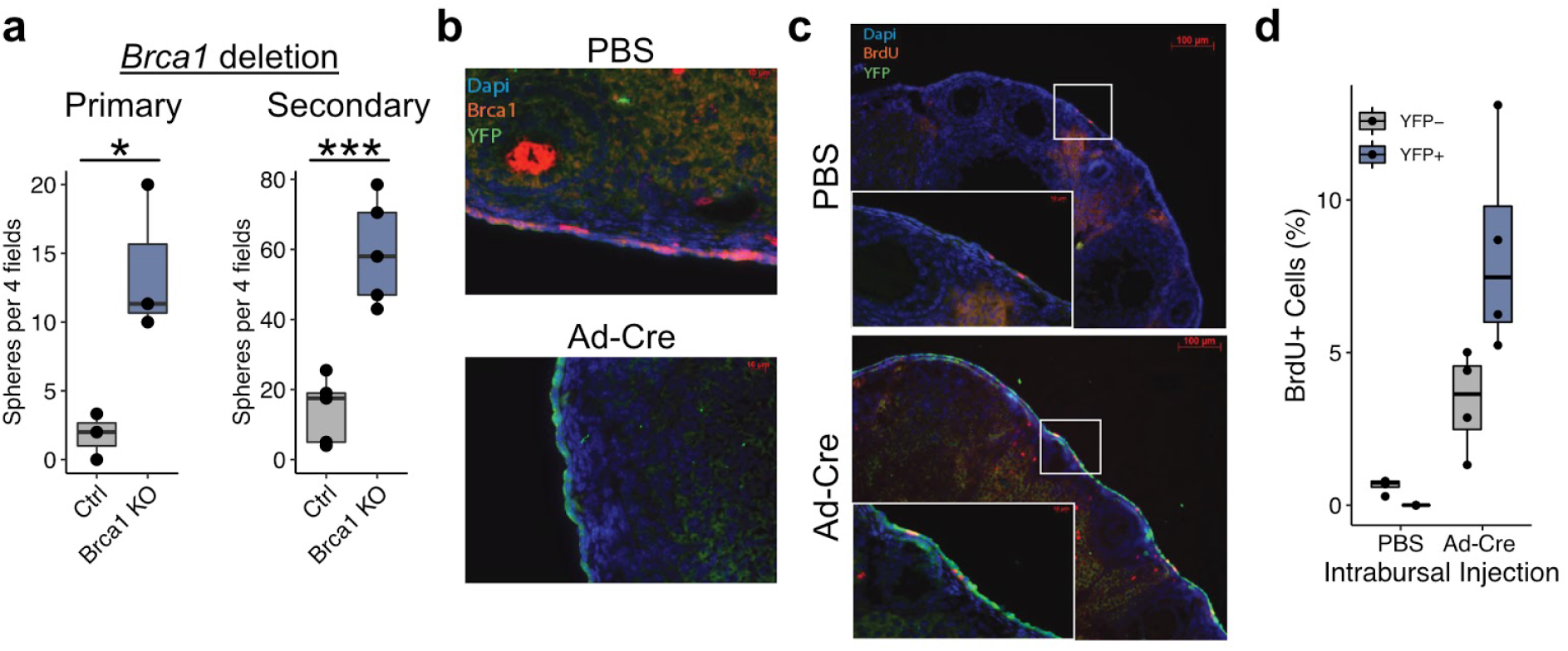
*Brca1* deletion *in vivo* promotes increased label retention. **a.** Primary (left) and secondary (right) sphere forming capacity of mOSE cells following *Brca1* deletion by infection with Ad-Cre (Brca1 KO). Cells infected with Ad-GFP were used as a control (Ctrl). Data points represent the average number of spheres per 4 fields of view for each replicate. **b.** Immunohistochemical staining of ovaries following intrabursal injection of PBS or Ad-Cre. Staining shows BRCA1 (red) and the YFP (green) reporter activated upon delivery of Cre recombinase. Nuclei are stained with DAPI (blue). Scale bar = 10μm. **c.** BrdU label retention (red) and YFP (green) signal in ovaries following intrabursal injection of either PBS or Ad-Cre. Scale bar = 100μm. **d.** Quantification of both BrdU+ cells in ovaries. All boxplots show median value (horizontal black line), estimated 25th and 75th percentiles, and whiskers represent 1.5 times the interquartile range.

To assess if BRCA1 loss enhances stemness phenotypes *in vivo*, we crossed the *Brca1^tm1Brn^* mice with *B6.129X1-Gt(ROSA)26Sor^tm1(EYFP)Cos^/J* mice to generate a *Brca1^fl/fl^YFP* mouse line, allowing us to track *Brca1*-null cells following exposure to Ad-Cre. These mice were injected intrabursally (IB) with Ad-Cre or PBS, and injected intraperitoneally (IP) with bromodeoxyuridine (BrdU). Ovaries were collected after a 30 day chase period and assessed for retention of the BrdU label and activation of the YFP reporter. Ad-Cre injection IB in *Brca1^fl/fl^YFP* mice showed successful activation of the YFP reporter, compared to the PBS injection (**Fig. 5b**). When combining the IB injections with an IP BrdU injection, Ad-Cre treatment increased the number of label-retaining OSE cells (**Fig. 5c,d**). This increased label retention further supports that BRCA1 loss can expand or induce populations of stem-like cells in the OSE.

To determine if BRCA1 loss results in similar expression patterns to other conditions associated with stemness, we performed RNA-seq on mOSE cells isolated from these mice infected with Ad-Cre or Ad-GFP. BRCA1 loss resulted in a large shift in gene expression, with 1499 significantly upregulated genes and 1881 downregulated (*p* < 0.05, absolute log fold change > 0.5; **Fig. 6a**; **Supplemental Data 7**). In mammary epithelium, the induction of EMT through *Brca1* deletion was presumed to be due to loss of BRCA1-mediated repression on the promoter of the EMT transcription factor *Twist1*. In contrast, we found that *Twist1* was approximately 8-fold lower in Brca1-null mOSE cells (**Fig. 6a**). There were also no EMT-associated gene sets enriched in upregulated genes. Rather, upregulated genes were largely enriched for gene sets associated with cell membrane transporters and downregulated genes were associated with cell cycle, oxidative phosphorylation, and DNA repair (**Fig. 6b; Supplemental Data 8**). *Brca1* deletion did not result in other features of stemness we observed in previous conditions, including activation of ERK and NFkB, and repression of interferon alpha response genes (**Fig. 6c**). *Brca1*-null cells were associated with reduced PI3K signalling and higher levels of estrogen signalling (**Fig. 6c**). Estrogen has been linked to EMT and stemness in other cell types^29^, but this is presumed to be through crosstalk, activating growth factor signalling pathways, which we do not see in *Brca1*-null OSE cells.

**Figure 6.**
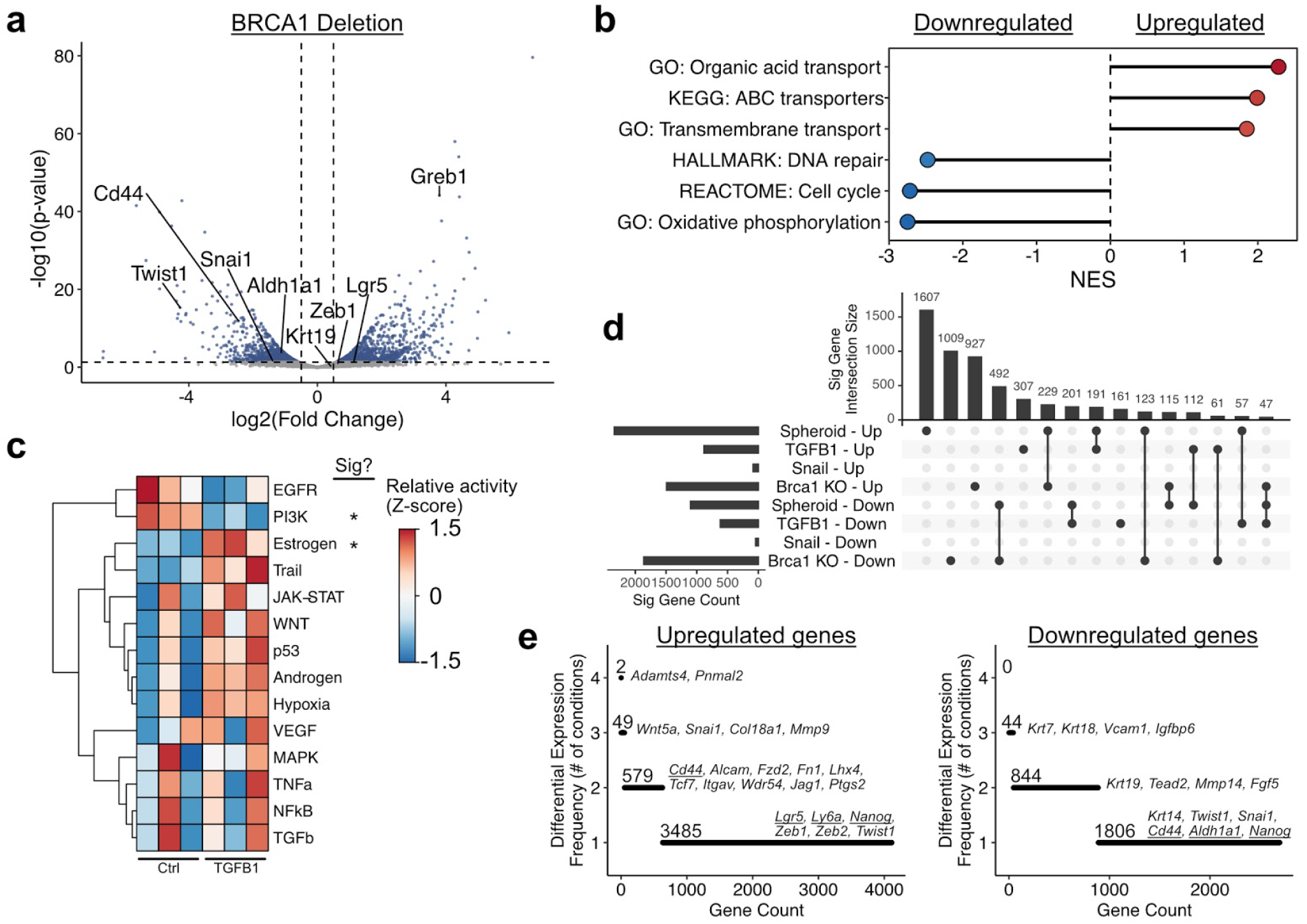
Stemness phenotypes are transcriptionally diverse. **a.** Plot showing the distribution of differentially expressed genes following *Brca1* deletion by infection with Ad-Cre. Each point corresponds to a single gene. Selected genes related to stemness and/or the EMT are highlighted on the plot. Dashed lines correspond to significance criteria (absolute log fold change > 0.5, *p*<0.05). **b.** GSEA results for selected gene sets enriched in differentially expressed genes following *Brca1* deletion. All gene sets are significantly enriched (*p* <0.05) and NES values are shown. **c.** Inferred pathway activity in control and Snail-overexpressing mOSE cells. *P*-values were computed from the *t* statistic of a linear regression model and were adjusted using the Benjamini-Hochberg FDR method. **d.** UpSet plot showing overlaps in differentially expressed genes between all conditions assessed ranked by the condition/overlap with the largest number of genes. **e.** Plots showing the number of assessed conditions that genes are either activated or repressed in. Selected genes are listed and putative stemness markers are underlined.

Comparing the expression profiles of each condition associated with stemness phenotypes in this study, we find minimal overlap in the specific genes activated or repressed in each (**Fig. 6d**). Ranking genes by the number of conditions they are activated or repressed in, we found that *Adamts4* and *Pnmal2* are the only genes upregulated in all four conditions (**Fig. 6e**). *Adamts4* has been linked to stemness in uveal melanoma through modulating crosstalk between the cells and their adjacent ECM^30^. While this may be relevant here, conserved downstream signals promoting stemness remain elusive. We note high frequency of EMT-associated changes, including activation of Snail, various collagens, and repression of cytokeratins (**Fig. 6e**). Notably, however, specific EMT transcription factors and putative OSE stemness markers (*Lgr5, Aldh1a1, Ly6a, Nanog*, and *Cd44*) are only activated in 1-2 conditions, and are even repressed in some conditions (**Fig. 6e**).

## Discussion

Several studies have reported putative stem cell populations in the OSE, but the relationships between these populations are unclear. The ability of differentiated epithelial cells to dedifferentiate and fulfil functional roles of stem cells has now been observed in several tissues, suggesting that static stem cell populations may not be required to maintain all tissues. In this study, we have further explored the ability of OSE cells to acquire features of stemness and have demonstrated it can be promoted by a variety of conditions. It may be expected that a common gene expression program would underlie the stemness phenotype, but we demonstrate that expression profiles are context-specific.

While transcriptional responses were variable, several patterns were recurrent across multiple conditions. We observed that induction of an EMT with TGFB1 treatment or Snail overexpression could promote stemness in OSE, but profiles of spheroids without cytokine treatment also showed EMT activation, which has also been seen with ovarian cancer cells^31^. As spheroid culture has been shown to enrich for cells with stem cell properties, these findings suggest that intrinsic stemness—independent of exogenous treatments—may be associated with a more mesenchymal phenotype. The relationship between the EMT and stemness is well documented^12,32^, but there is growing evidence that stemness and EMT are not inextricably linked. For example, the EMT-promoted transcription factor PRRX1 suppresses stemness in breast cancer cells^33^. Further, transcriptional dynamics of the EMT have been shown to be highly context-specific, which explains why it does not consistently promote stemness^25^. Just as the EMT can occur without promoting stemness, we have shown that deletion of *Brca1* promotes stemness in OSE without activating any EMT-associated expression, including *Twist1* activation, which had been shown to drive stemness following *Brca1* deletion in mammary epithelial cells^28^. Instead, *Brca1* loss caused many changes in cell membrane transport and metabolic genes. While the mechanism of induced stemness following *Brca1* loss is unclear, this provides strong evidence that stemness is not dependent on a mesenchymal expression profile and is perhaps as context-specific as the EMT response^25^.

Several alterations in signalling pathway activity were also common across conditions. TGFB1 treatment and spheroids were associated with higher levels of ERK activity, which have both been linked to stemness in epithelial^34^ and carcinoma cells^35^. In EGF-free media, paracrine/autocrine signalling is established, maintaining ERK activity in stem cell populations of intestinal organoids^35^. While these mechanisms may contribute to stemness in OSE spheroids or those treated with TGFB1, increased ERK activity was not enhanced following Snail overexpression or *Brca1* loss. Similarly, a gene set comprising interferon alpha response genes was downregulated following spheroid culture, TGFB1 treatment, and Snail overexpression. While it is unlikely that interferon alpha itself was present, it is possible that various signalling pathways may affect common target genes. Consistent with this, disruption of type 1 interferon signalling promotes stemness in breast cancer cells^36^. None of these patterns, however, are consistent across all conditions, further supporting that mechanisms promoting stemness may vary considerably depending on environmental conditions (eg. ovulatory wound repair, tissue expansion during folliculogenesis, natural cell turnover).

While we have relied heavily on *in vitro* models here, this has enabled us to explore the ability of OSE cells to acquire stemness following various experimental perturbations. This suggests that differentiated epithelial cells may be capable of self-regulating tissue maintenance in response to environmental cues, such as tissue damage. The expression profiles of this emergent stemness may be variable, depending on the specific properties of the cells’ microenvironment. This model is particularly interesting because it is a stark contrast to how stem cells and differentiation hierarchies have been viewed for the last several decades. The OSE is a promising tissue to explore this further as it undergoes regular rupture and repair throughout reproductive cycles, and is a simple tissue comprising a single cell type, which may be the most likely to exhibit this behavior. Designing strategies to monitor stemness dynamics *in vivo* will be critical to understand these behaviours in a normal physiological context.

## Methods

### OSE cell isolation and culture

The isolation and culture of mOSE cells was done as previously described^10^, in accordance with the guidelines of the Canadian Council on Animal Care and under a protocol approved by the University of Ottawa Animal Care Committee. Briefly, ovaries from randomly cycling female mice (FVB/N, 6 weeks old) were collected and incubated in 0.25% Trypsin/PBS (Invitrogen) (37 °C, 5% CO_2_, 30 min) to facilitate OSE removal. mOSE cells were isolated by centrifugation and plated onto tissue culture plates (Corning) in mOSE media [a-Minimum Essential Medium (Corning) supplemented with 4% FBS, 0.01 mg/mL insulin-transferrin-sodium-selenite solution (ITSS; Roche), and 2 μg/mL EGF (R&D Systems)]. hOSE cells were isolated and cultured as previously described ^37^, with patient consent and under a protocol approved by the Ottawa Health Science Network Research Ethics Board (Protocol #1999540). Briefly, ovaries from 5 different women were collected during surgery for reasons other than ovarian pathology. Using a scalpel, hOSE cells were scraped from the ovarian surface and isolated by centrifugation in hOSE media (Wisent Bioproducts) supplemented with 10% FBS. All mouse and human OSE cells were passaged 2-3 times prior to experimental use and experiments were conducted with cells of a passage number less than 25.

### Quantitative reverse transcription polymerase chain reaction (RT-PCR)

The RNeasy Mini Kit (Qiagen) was used to extract RNA and the OneStep RT-PCR Kit (Qiagen) was used to synthesize cDNA. Quantitative PCR was done using the ABI 7500 FAST qRT-PCR machine (Applied Biosystems) using the Taqman gene expression (Life Technologies) and SsoFast gene expression (Bio-rad) assays utilizing *Tbp* as an endogenous control. Primer sequences are listed in Supplemental Table 1. RQ (relative quantity) was determined using the cycling threshold for the gene of interest in control or untreated samples compared to the cycling threshold in experimental samples, calculated using the Applied Biosystems 7500 FAST v2.3 software.

### Western blot

M-PER mammalian protein extraction reagent (GE Healthcare) was used to extract protein from mOSE cells and run on NuPAGE 4-12% Bis-Tris gradient gels (Life Technologies).

Polyvinylidene difluoride membranes were used to transfer protein samples. Membranes were blocked in 5% non-fat milk prior to antibody incubation. Antibody conditions are described in Supplemental Table 2. Western blots were developed using Clarity™ Western ECL Substrate (Bio-Rad) and the FluorChem FC2 imaging system (Alpha Innotech).

### Stem cell PCR array

mOSE cells (1×10^6 cells) were plated 24 hr prior to treatment with TGFB1 (10 ng/mL, R&D Systems). RNA was collected 7 days post-TGFB1 treatment (RNAeasy Kit, Qiagen). cDNA synthesis was performed using RT^2 First Strand Kit (Qiagen) and run on the RT^2 First Strand Kit (Stem cell PCR array) (Qiagen). The array was run in triplicate (N=3) and analyzed using the DataAnalysis Excel platform provided with the array kits.

### Snail-overexpressing mOSE cells

mOSE cells stably expressing reverse tetracycline-controlled transactivator (rtTA) protein were transduced with a lentiviral construct (pWPI) expressing the murine *Snai1* or *eGFP* under the control of a doxycycline-inducible promoter and the hygromycin resistance gene under the control of the *PGK* promoter. Transduced cells were selected for resistance to Hygromycin B. 200ng/mL of doxycycline was added to cultures for 4 days prior to all experiments overexpressing Snail.

### Primary sphere-forming assays

For free-floating spheres, mOSE cells were cultured in stem cell media [Dulbecco’s Modified Eagle’s Medium: Nutrient Mixture F-12 (Sigma) supplemented with 1 X B27 supplement (Invitrogen), 0.02 μg/mL EGF (R&D Systems), 0.04 μg/mL fibroblast growth factor (FGF; R&D Systems), 4 μg/mL heparin (Sigma) and 0.01 mg/mL ITSS (Roche), and 2 μg/mL EGF (R&D Systems)] at 5×10^4 cells/mL in non-adherent 24-well culture plates (Corning) and incubated at 37 °C, 5% CO_2_ for 14 days. Spheres were quantified using ImageJ using a pixel cutoff of >1000 pixels and a circularity limit of 0.5-1.0. For spheres cultured in methylcellulose, mOSE cells were placed in a 1:1 mixture of methylcellulose and stem cell media at 5×10^4 cells/mL in 24-well culture plates (Corning), and incubated at 37 °C, 5% CO_2_ for 28 days.

Methylcellulose-embedded spheres were quantified using ImageJ using a pixel cutoff of >500 pixels and a circularity limit of 0.5-1.0. For each experiment, a minimum of 3 replicates were performed, each replicate was performed in three independent wells, spheres were counted in 4 fields per well, and the average count was reported.

### Secondary sphere-forming assay

Primary free-floating mOSE spheres were collected and washed in PBS. Spheres were dissociated by first incubating in trypsin/PBS (Invitrogen) at 37 °C for 10 min, then by passing cells through a 25 gauge needle to obtain a single cell suspension. Single cell suspension was verified using phase contrast microscopy. Cells were washed in PBS, counted using a hemocytometer and plated in stem cell media at 5×10^4 cells/mL. Cells were incubated in non-adherent 24-well culture plates (Corning) at 37 °C, 5% CO_2_ for 14 days. Spheres were quantified using ImageJ using a pixel cutoff of >500 pixels and a circularity limit of 0.5-1.0.

### *Brca1* deletion in mOSE cells

mOSE cells were isolated from homozygous *Brca1^tm1Brn^* mice as described above and then infected with Ad-Cre to achieve *Brca1* knockout. Ad-GFP was used as a control. Cells were cultured for 1 week after infection prior to experimental use.

### BrdU pulse-chase

*Brca1^tm1Brn^* mice were bred to *B6.129X1-Gt(ROSA)26Sor^tm1(EYFP)Cos^/J* mice to produce *Brca1^fl/fl^YFP* mice. *Brca1^fl/fl^YFP* mice were injected IB with Ad-Cre (8×10^7^ PFU) or PBS on day 1 and injected IP with BrdU (0.25 mg daily) on days 7-10. Ovaries were collected on day 40 and frozen in Optimal Cutting Temperature Compound.

### BrdU Immunofluorescence

Frozen sections (5 μm) were fixed using formalin-vapor fixation overnight at −20 °C. Samples were then hydrated in PBS and antigen retrieval performed using an antigen unmasking solution (ph 6.0, Vector) in a steam chamber (Hamilton Beach). Slides were then washed in PBS and blocked with 5% goat serum for 1 hr at room temperature. Primary antibodies against BRCA1 (1:200, H-300, rabbit), GFP (1:1000, ab13970, chicken), and BrdU (1:200, ab6326, rat) were added and incubated overnight at 4 °C. Following a PBS wash, species-appropriate secondary antibodies (1:250, Alexafluor 594 nm or 488 nm) were incubated for 1 hr at room temperature. Slides underwent a final PBS wash and were mounted using Prolong Gold with DAPI (ThermoFisher). Positive cells were counted manually.

### RNA-seq sample preparation

For TGFB1 treatment of monolayer cultures, mOSE cells were plated 24hrs prior to the addition of TGFB1 (10ng/mL, R&D Systems) and cells were collected after 4 days of treatment. For inducible *Snail* expression and *Brca1* deletion, cells were plated for 24hr prior to the addition of doxycycline (200 ng/mL, Sigma), and RNA was collected 4 days later. For sphere-forming conditions, mOSE cells (1×10^6) were first plated as monolayer cultures 24 hr prior to treatment with TGFB1 (10 ng/mL, R&D Systems). Four days after the addition of TGFB1, mOSE cells were then plated in free-floating sphere-forming conditions. Cells were maintained in sphere-forming cultures for 2 weeks prior to RNA collection (RNAeasy Kit, Qiagen). TGFB1 was replenished when placing mOSE cells in sphere-forming conditions.

### Library preparation and sequencing

Total RNA was quantified using a NanoDrop Spectrophotometer ND-1000 (NanoDrop Technologies, Inc.) and its integrity was assessed on a 2100 Bioanalyzer (Agilent Technologies). Libraries were generated from 250 ng of total RNA as follows: mRNA enrichment was performed using the NEBNext Poly(A) Magnetic Isolation Module (New England BioLabs). cDNA synthesis was achieved with the NEBNext RNA First Strand Synthesis and NEBNext Ultra Directional RNA Second Strand Synthesis Modules (New England BioLabs). The remaining steps of library preparation were done using the NEBNext Ultra II DNA Library Prep Kit for Illumina (New England BioLabs). Adapters and PCR primers were purchased from New England BioLabs. Libraries were quantified using the Quant-iT™ PicoGreen^®^ dsDNA Assay Kit (Life Technologies) and the Kapa Illumina GA with Revised Primers-SYBR Fast Universal kit (Kapa Biosystems). Average size fragment was determined using a LabChip GX (PerkinElmer) instrument.

The libraries were normalized, denatured in 0.05 N NaOH and then diluted to 200 pM and neutralized using HT1 buffer. ExAMP was added to the mix and the clustering was done on an Illumina cBot and the flowcell was run on a HiSeq 4000 for 2×100 cycles (paired-end mode) following the manufacturer’s instructions. A phiX library was used as a control and mixed with libraries at 1% level. The Illumina control software was HCS HD 3.4.0.38 and the real-time analysis program was RTA v. 2.7.7. The program bcl2fastq2 v2.18 was then used to demultiplex samples and generate fastq reads.

### RNA-seq processing and differential expression

Transcript quantification for each sample was performed using Kallisto (v0.45.0)^38^ with the GRCm38 transcriptome reference and the −b 50 bootstrap option. The R package Sleuth (v0.30.0) ^39^ was then used to construct general linear models for the log-transformed expression of each gene across experimental conditions. Wald’s test was used to test for significant variables for each gene and the resultant p-values were adjusted to q-values using the Benjamini-Hochberg false discovery rate method. Significant genes were defined as genes with a q-value < 0.05. An effect size (beta coefficient of the regression model) cutoff of >0.5 or <-0.5 was also used for each data set.

### Gene set enrichment analysis and pathway activity inference

GSEA was performed with the R package fgsea (v1.13.5)^40^. GO terms, KEGG pathways, Reactome pathways, and Hallmark genesets were collected from the Molecular Signatures Database (MSigDB)^22,23^ and used to query differential expression results ranked by fold change. All gene sets discussed in the manuscript have a significant enrichment (Benjamini-Hochberg adjust p-value <0.05). For pathway activity inference, we used the R package PROGENy (v1.9.6)^41^. Pathway activity was compared between experimental conditions using a simple linear model and p-values were adjusted using the Benjamini-Hochberg false detection rate method.

### CD44 cell sorting

mOSE cells were treated with TGFB1 (10 ng/mL, 2 days) prior to collecting cells for FACS. Cells (1×10^7) were trypsinized and a single-cell suspension was made using a 40 μm cell strainer. Cells were labelled and sorted as previously described ^18^. Briefly, cells were resuspended in a flow buffer (4% FBS in PBS) and incubated with anti-CD44 conjugated to allophycocyanin (1:5000; eBioscience, San Diego, CA) for 15 min at 4 °C. Unbound antibody was removed with washing buffer and the fraction of cells with surface protein labeled with CD44 antibody was determined using a MoFlo cell sorter (Dako Cytomation).

### Data availability

Raw sequencing files have been deposited and are available along with processed transcript quantifications at GSE122875.

## Acknowledgments

We thank the tissue donors for making this research possible. We also thank Dr. Ken Garson for generating mOSE cells with inducible *Snail* and inducible *GFP* expression. We wish to acknowledge the contribution of staff of the McGill University and Génome Québec Innovation Centre (Montreal, QC) for performing library preparation and sequencing associated with RNA-seq experiments. We also wish to acknowledge StemCore Laboratories (Ottawa, ON) for performing the FACS of CD44-positive mOSE cells. This work was supported by grants from the Canadian Institutes of Health Research and the National Science and Engineering Research Council (BCV). L.E.C. and L.F.G. were supported by Ontario Graduate Scholarships, D.P.C. by a Frederick Banting and Charles Best Doctoral Award (CIHR), and C.W.M. by the Vanier Canada Graduate Scholarship.

## Author Contributions

L.E.C. and B.C.V. conceived the study. L.E.C., D.P.C., and B.C.V. interpreted results and wrote the manuscript. L.E.C., L.F.G., O.C., H.A.D., and T.D. performed cell culture experiments, qPCR analysis, and western blots. O.C. derived mOSE and hOSE cultures. C.W.M. performed mouse experiments and immunofluorescence. D.P.C. and L.E.C. performed all computational analysis.

## Competing Interests

The authors declare no competing interests.

---

**Supplemental Figure 1.**
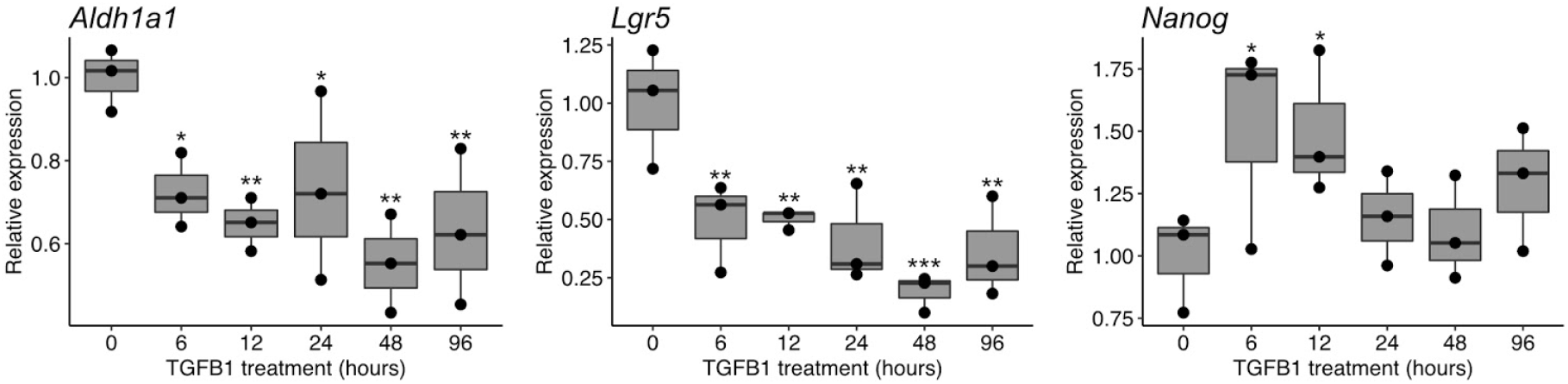
Expression of stemness markers through a time course of TGFB1 treatment. qPCR results showing expression of stemness genes following varying lengths of TGFB1 treatment. Boxplots show median value (horizontal black line), estimated 25th and 75th percentiles, and whiskers represent 1.5 times the interquartile range. *P*-values were computed from the *t* statistic of a linear regression model. * *p*<0.05, ** *p* <0.01, *** *p* <0.001.

**Supplemental Figure 2.**
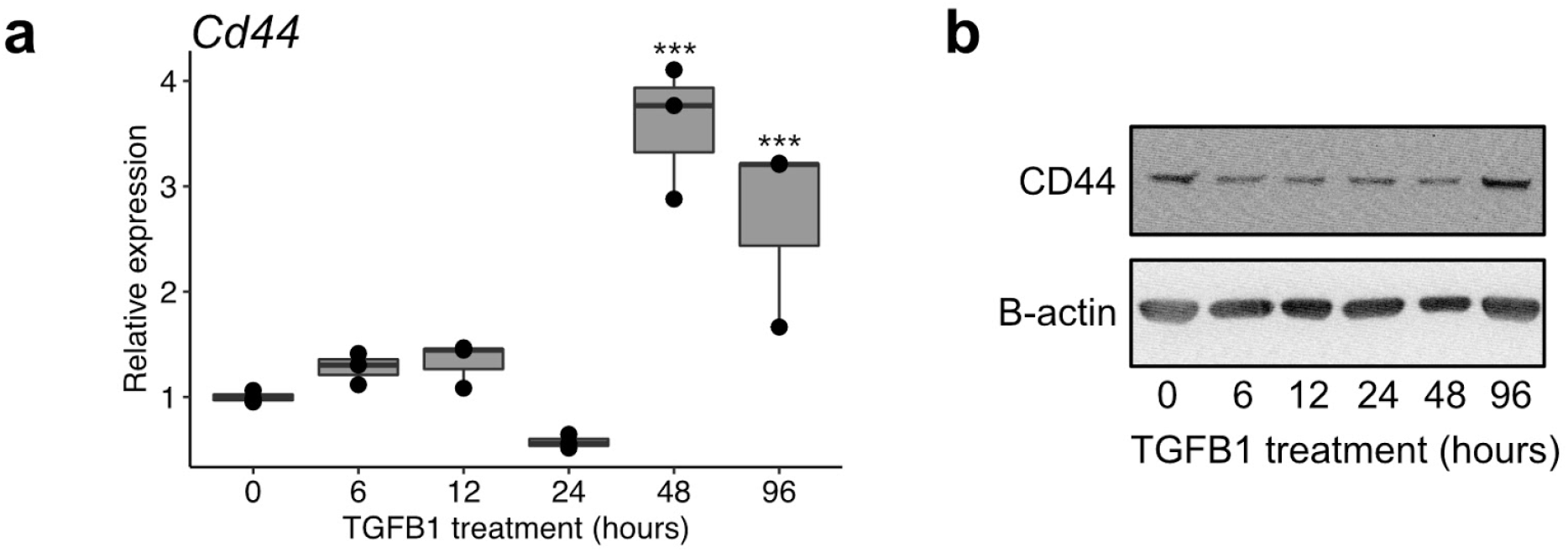
CD44 increases following TGFB1 treatment. **a.** qPCR results showing expression of *Cd44* following varying periods of TGFB1 treatment. Boxplots show median value (horizontal black line), estimated 25th and 75th percentiles, and whiskers represent 1.5 times the interquartile range. *P*-values were computed from the *t* statistic of a linear regression model. * *p* <0.05, ** *p* <0.01, *** *p* <0.001. **b.** Representative western blot of CD44 and B-actin following following varying periods of TGFB1 treatment. Densitometric quantifications of three blots are included in Figure 1e.

**Supplemental Figure 3.**
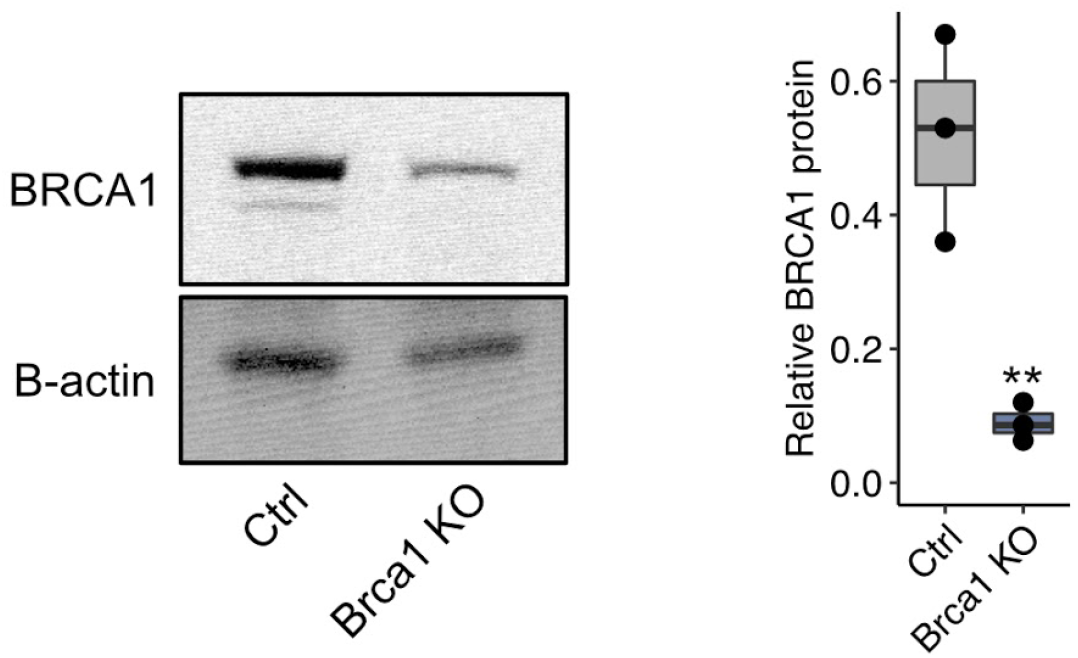
*Brca1* is efficiently deleted following the expression of Cre recombinase. **a.** Western blot of BRCA1 and B-actin of *Brca1^fl/fl^* mOSE cells following infection with either Ad-GFP (Ctrl) or Ad-Cre (Brca1 KO). **b.** Boxplot of relative BRCA1 protein quantifications from pixel densitometry. N=3 independent infections with Ad-GFP or Ad-Cre. Boxplots show median value (horizontal black line), estimated 25th and 75th percentiles, and whiskers represent 1.5 times the interquartile range. *P*-values were computed from the *t* statistic of a linear regression model. ** *p* <0.01.

## Notes

### Competing Interest Statement

The authors have declared no competing interest.

